# Structural evolution of a fungal cell wall protein family for β-glucan-binding and cell separation

**DOI:** 10.1101/2025.11.20.689653

**Authors:** Philipp Schöppner, Vitali Weitzel, Maik Veelders, Lukas Korf, Jonas Andräs, Katharina Wolf, Stefan Brückner, Lars-Oliver Essen, Hans-Ulrich Mösch

## Abstract

In fungi, the continuous biosynthesis and remodeling of the cell wall are crucial for growth, division, and development. A hallmark of fungal cell walls is their layered structure, which includes several gel-like carbohydrate polymers, such as β-glucans, and a large number of associated cell wall proteins. The fungal-specific family of SUN domain proteins has been implicated in cell wall remodeling and cell separation, but detailed structure-based analyses revealing precise molecular functions have been lacking until now. In this study, we determined high-resolution crystal structures of the SUN domains from two paralogs of the SUN family in budding yeast. We find that their bilobal architecture consists of a sushi-like domain and an intimately associated thaumatin-like domain, which together form a highly conserved canyon fitted to accommodate both single- and triple-helical β-glucan polymers. Within this canyon, we identify twelve conserved polar residues that are crucial for the function of SUN domains in mediating cell separation. We further demonstrate that SUN domains are functionally interchangeable between paralogs in budding yeast, as well as between orthologs from budding yeast and phylogenetically distant fission yeast or filamentous fungi. We conclude that the fungal SUN domain family represents a unique class of β-1,3-glucan-binding proteins involved in cell wall remodeling and separation, whose successful evolution was enabled by the fusion of ancestral sushi- and thaumatin-like domains.

**Importance:** Fungal cell walls are dynamic extracellular structures essential for growth and morphogenesis, making them prime targets for antifungal drugs and the host immune system. Although many protein families involved in the synthesis, crosslinking, and degradation of cell wall polymers are known, the molecular functions and structural evolution of most cell wall proteins remain poorly understood. Our in-depth structural, functional, and phylogenetic analysis of the fungal SUN domain protein family sheds light on a central question: how specific protein families have evolved structurally to enable dynamic cell wall remodeling during growth and division. Moreover, this work identifies precise structural targets within the fungal cell wall that could guide the development of novel diagnostics and therapeutics against life-threatening fungal infections.

## Introduction

Fungal cell walls are highly dynamic extracellular organelles with a thickness of 50 to 250 nm, which confer mechanical strength, morphological attributes, environmental protection and adhesive properties (1–3). Comprehensive genome-wide studies in yeast and filamentous fungi have shown that approximately 20 % of their genes are involved in cell wall structure, function, biosynthesis and dynamics, emphasizing the fundamental importance of fungal cell walls (4, 5). To fulfill their functions, fungal cell walls have a layered architecture and mainly consist of chitin, β-glucans, mannans and associated cell wall proteins (CWPs) (2, 6–8). In yeasts and filamentous fungi of the phylum of Ascomycota, the inner cell wall layer is an elastic and porous network, which mainly consists (65% to 90%) of long linear chains of β-1,3-glucans and a small fraction of variable β-1,6-linked glucan side chains (9), which can serve as covalent attachment sites for GPI-anchored CWPs (7, 10, 11). The majority of linear β-1,3-glucans adopt structurally well-defined single- or triple-helical conformations, whose fibrillar structures are stabilized by hydrogen bonds (12, 13). Known cross-linkers of the β-glucan network are CWPs of the Pir family (<u>P</u>roteins with internal repeats), which are O-ester-linked to glucan hydroxyl groups via deamidated glutamine side chains (14).

Fungal cell walls not only need to be rigid structures, but they also require continuous remodeling during cell growth, cell separation and morphological development (15, 16). For this purpose, fungal cells harbor a large number of secreted enzymes for synthesis, crosslinking and degradation of cell wall carbohydrate polymers (8, 17, 18). Prominent examples include Fks1 and Chs1 for β-1,3-glucan and chitin synthesis (19), respectively, Gas family proteins for crosslinking via β-1,3-glucanosyltransferase activity (20), Crh-type transglycosylases for chitin to β-1,6-glucan linking activity (21) and degrading enzymes such as chitinases and glucanases (22, 23). In addition, numerous genes have been identified that affect cell wall remodeling, including *SUN*, *SCW* and *CCW* genes from the budding *Saccharomyces cerevisiae* and corresponding orthologs in other fungi (8, 18). However, the precise molecular functions for many of the encoded proteins remain largely unknown.

The SUN family of proteins is widely distributed in the fungal phylum Ascomycota and characterized by the presence of the well-conserved SUN domain (INTERPRO IPR005556). Currently, the family contains roughly two thousand members, most of which consists of a SUN domain, which is fused to a N-terminal low-complexity region rich in serine and threonine residues. SUN family protein have further been subdivided into two groups based on an additional C-terminal region, which is variable in length and that is lacking in members of group I (Sun4-type), but can be found in group II (Adg3-type) members (24–26). Moreover, SUN family proteins are predicted to be secreted into the periplasmic space and to lack anchoring to the plasma membrane via transmembrane domains or glycosyl-phosphatidyl-inositol (GPI) anchors. So far, a few *SUN* genes have been functionally characterized by genetic approaches in a small number of Ascomycota, which cover the three subphyla Taphrinomycotina (fission yeasts), Saccharomycotina (budding yeasts) and Pezizomycotina (filamentous fungi) (27). In the budding yeast *S. cerevisiae*, which belongs to the subphylum of Saccharomycotina, the SUN family includes five genes, *ScSUN4*, *ScSIM1*, *ScUTH1*, *ScNCA3* and *ScADG3* (*YMR244W*). While *ScSUN4* and *ScUTH1* have been attributed a role in cell wall remodeling, as exemplified by the failure of corresponding mutants to confer efficient cell separation (24, 28), no clear cell wall function has been attributed to other SUN family members in *S. cerevisiae*. In the human pathogen *Candida albicans*, which forms part of the Saccharomycotina and contains three different *SUN* family genes, strains lacking *CaSUN41* display reduced cell separation and hyphal elongation (25, 29, 30). In the fission yeast *Schizosaccharomyces pombe*, which belongs to the Taphrinomycotina and harbors three *SUN* family genes, strains lacking the *SpPSU1* gene show severe cell wall damages (31). In *Botrytis cinerea*, a filamentous plant pathogen of the Pezizomycotina harboring two SUN family genes, *BcSUN1* has been shown to be involved in maintaining the structure of the cell wall and/or extracellular matrix (32). Finally, in the human pathogen *Aspergillus fumigatus*, a filamentous fungus that forms part of the Pezizomycotina and contains two SUN domain genes, mutation of *AfSUN1* causes severe damages in the hyphal cell wall (26). In addition, a biochemical function has been described for AfSun1, which exhibits a weak exo-β-1,3-glucanase activity *in vitro* (26). Nonetheless, the precise molecular function of most SUN domain proteins remains elusive, and it is currently not known, how functional diversity of different paralogs has evolved.

In this work, we present high-resolution 3D-structures of the SUN domains of *S. cerevisiae* ScSun4 and ScSim1, which encompass an N-terminal sushi-like domain followed by a thaumatin fold. Together, the two domains form a canyon-like shape, whose dimensions are complementary to the structures of single and triple-helical β-1,3-glucans. Mutational analysis identifies twelve conserved residues within this canyon, which significantly contribute to ScSun4-mediated control of cell separation. Remarkably, SUN domains are functionally transferable between paralogs as well as between orthologs of phylogenetically very distant fungi. We therefore propose that the fungal-specific SUN domain family represents a unique class of single and triple-helical β-glucan binding proteins, which acquired their function through fusion of ancestral sushi- and thaumatin-like domains.

## Results

### Phylogenetic analysis of the fungal SUN domain family

To obtain a general overview of the fungal SUN domain family, we subjected 1602 SUN domains present in the INTERPRO database (IPR005556) to a comprehensive comparative analysis by employing the Enzyme Similarity Tool (EFI-EST) (33). The resulting protein sequence similarity network (SSN) visualizes the broad relationships (Figure 1A) and supports the previous classification into two major groups (24, 26), which differ by the absence (group I) or presence (group II), respectively, of a serine-threonine-rich (STR) domain of variable length C-terminal to the SUN domain. Overall, a total of 982 SUN domains (61.3 %) can be attributed to group I, whereas 620 domains (38.7 %) belong to group II, respectively. Our analysis also reveals, that the absence or presence of an additional transmembrane domain at the C-terminus of group II proteins does not correlate with a clear SSN-based subdivision of their SUN domains. It is further interesting to note, that >97 % of the fungal SUN domain proteins present in the INTERPRO database can be attributed to the fungal phylum Ascomycota, whereas SUN domains are absent in Basidiomycota. We further analyzed the SUN domains of 77 selected members from the phylum Ascomycota covering group I and group II by calculating a phylogenetic ML-tree, that is based on a T-Coffee alignment (Figure 1B). This analysis essentially reflects the division of SUN domains obtained by the SSN approach into group I and group II. In addition to the SSN analysis, the phylogenetic ML-tree analysis results in a clear subdivision of group I members into SUN domains present in filamentous fungi from the subphylum Pezizomycotina or in yeast-like fungi from the subphyla Saccharomycotina and Taphrinomycotina, respectively. Further sequence analysis of 20 selected SUN domains from this group of proteins shows that they share the well-conserved Cx_5_Cx_3_Cx_24-27_C cysteine-pattern (Figure S1A), although pairwise sequence identities can be as low as 26 % (Figures S1B and S1C).

**Figure 1.**
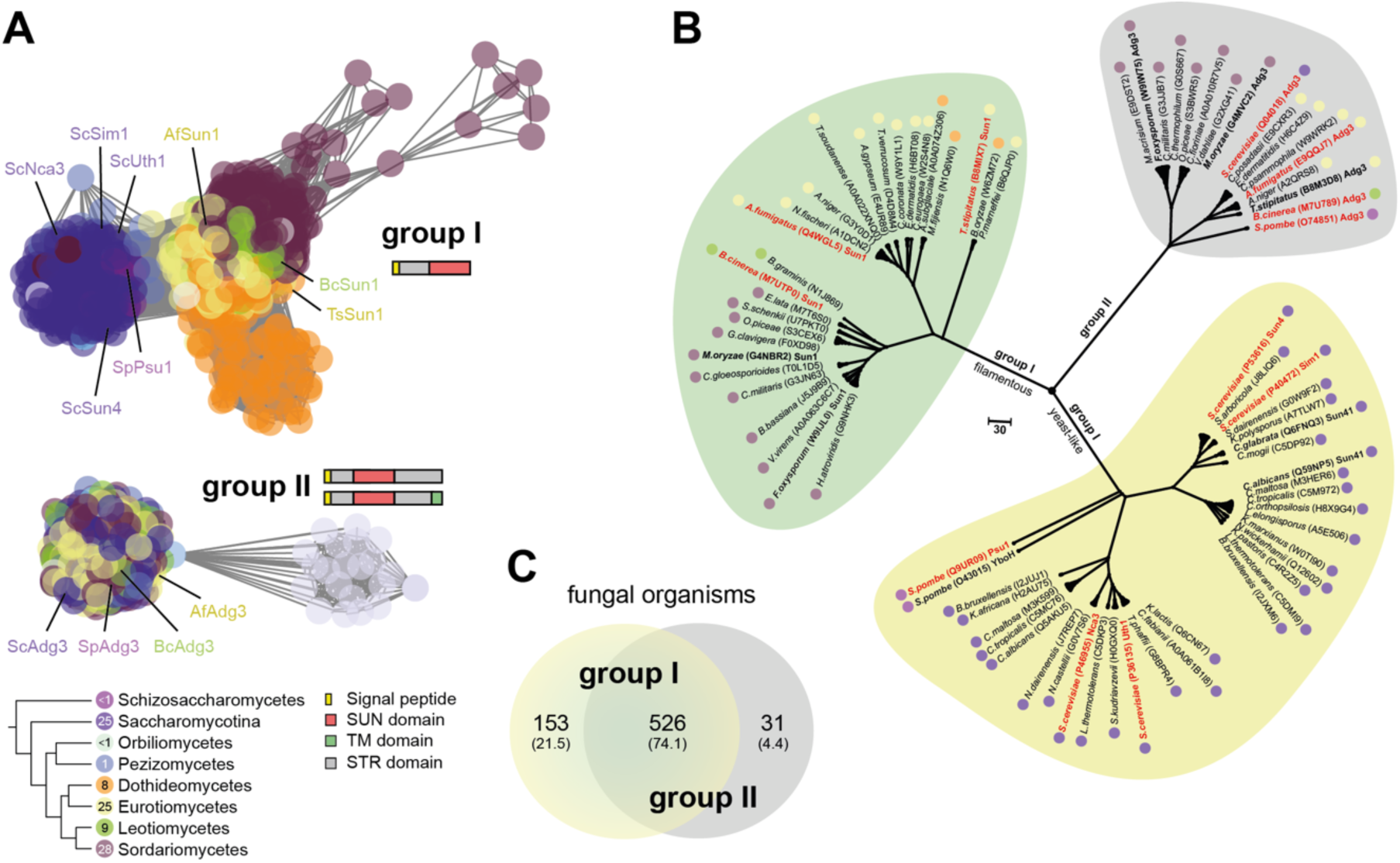
Phylogenetic analysis of the fungal SUN domain family. A) Sequence-similarity network (SSN; E-value cut-off 10^-60^) (33) of 1602 SUN domains from the fungal SUN domain family (IPR005556) reveals two major groups. More than 97 % of known SUN domain family members are found in the phylum Ascomycota as represented by colored dots according to the phylogeny code shown at the bottom left. Numbers in colored dots indicate the percentage of SUN domains present in the respective fungal taxa. Members of group I and group II, which were investigated *in vitro* and *in vivo*, are indicated by their name. The domain structures of group I and group II members are shown as colored diagrams according to the legend displayed at the bottom right. TM = transmembrane; STR = serine-threonine-rich. B) Phylogenetic ML-tree of 77 selected SUN domains based on a T-Coffee alignment. For each SUN domain the originating fungal species and protein (names according to the UniProt Knowledgebase) are shown together with colored dots indicating the originating fungal class (for color code see *A*). Branches of group I members from filamentous fungi are colored in green and from yeast-like fungi in yellow, respectively. The branch of group II members is colored in grey. Information shown in bold letters corresponds to the domains used for the multiple sequence alignment shown in Figure S1. Functionally investigated members shown in *A* are indicated in red color. A ruler for phylogenetic distances is shown near the center. C) VENN diagram showing the presence of group I (yellow circle) and group II (grey circle) SUN domains in 710 fungal organisms. Numbers indicate the organisms that contain only group I domains (153), only domains from group II (31), or domains from both groups (526), respectively. Corresponding percentage values are shown in parentheses.

We further determined the distribution patterns for a total of 1408 group I and group II SUN domains in 710 different fungal organisms with available genome sequences, covering a total of 607 different species (Figure S2A and Table S1). Here, we found that a large fraction of 526 organisms (74.1 %) contains domains from both groups, whereas 153 fungi (21.5 %) only contain group I domains, and 31 organisms (4.4 %) only carry domains from group II, respectively (Figure 1C). We further found that the number of paralogs ranged from two to six within organisms from yeast-like fungi of the subphyla Saccharomycotina and Taphrinomycotina (Figure S2B and Table S1). In addition, two or more group I proteins are present in most yeast-like organisms carrying three or more paralogs (Figures S2C and S2D; Table S1). In contrast most group II containing yeast-like fungi are restricted to a single group II paralog. Our analysis further shows that the number of paralogs is restricted to one to two in most filamentous fungi of the subphylum Pezizomycotina (Figures S2B). Among these filamentous fungi, most organisms of the classes of Sordariomycetes, Leotiomycetes, Eurotiomycetes and Pezizomycetes, respectively, contain a single paralog of each group I and group II (Figures S2C and S2D; Table S1). In contrast, Dothideomycetes are restricted to a single group I paralog and Orbiliomycetes to a single paralog of group II.

In summary, our analysis reveals that SUN domain proteins are highly restricted to the phylum of Ascomycota. Moreover, yeast-like organisms belonging to the subphyla of Saccharomycotina and Taphrinomycotina appear to contain increasing numbers of group I family members. In contrast, filamentous fungi of the subphylum Pezizomycotina are mostly restricted to 1 to 2 family members, whereby class-specific distribution patterns of group I and/or group II SUN domains seem to have evolved.

### High resolution structures of the SUN domains of *S. cerevisiae* Sun4 and Sim1

We next solved the crystal structures of fungal SUN domains, in order to obtain further insights into their molecular functions and to provide a basis for a structure-guided functional analysis. For this purpose, we first focused on Sun4 from *S. cerevisiae* (ScSun4), because this variant belongs to the best characterized SUN proteins so far and is well-suited for functional *in vivo* characterization. Analysis of full-length ScSun4 by the Regional Order Neural Network (RONN) (34) reveals a high probability of disorder for the N-terminal serine-threonine-rich region. Therefore, a shortened version of ScSun4 covering the conserved SUN domain (residues G147-N420) was produced by heterologous expression in *E. coli* and subsequent purification. After successful crystallization and X-ray analysis, the 3D structure of the SUN domain of ScSun4 could be determined at a resolution of 1.1 Å (Figure 2A left, Table S2).

**Figure 2.**
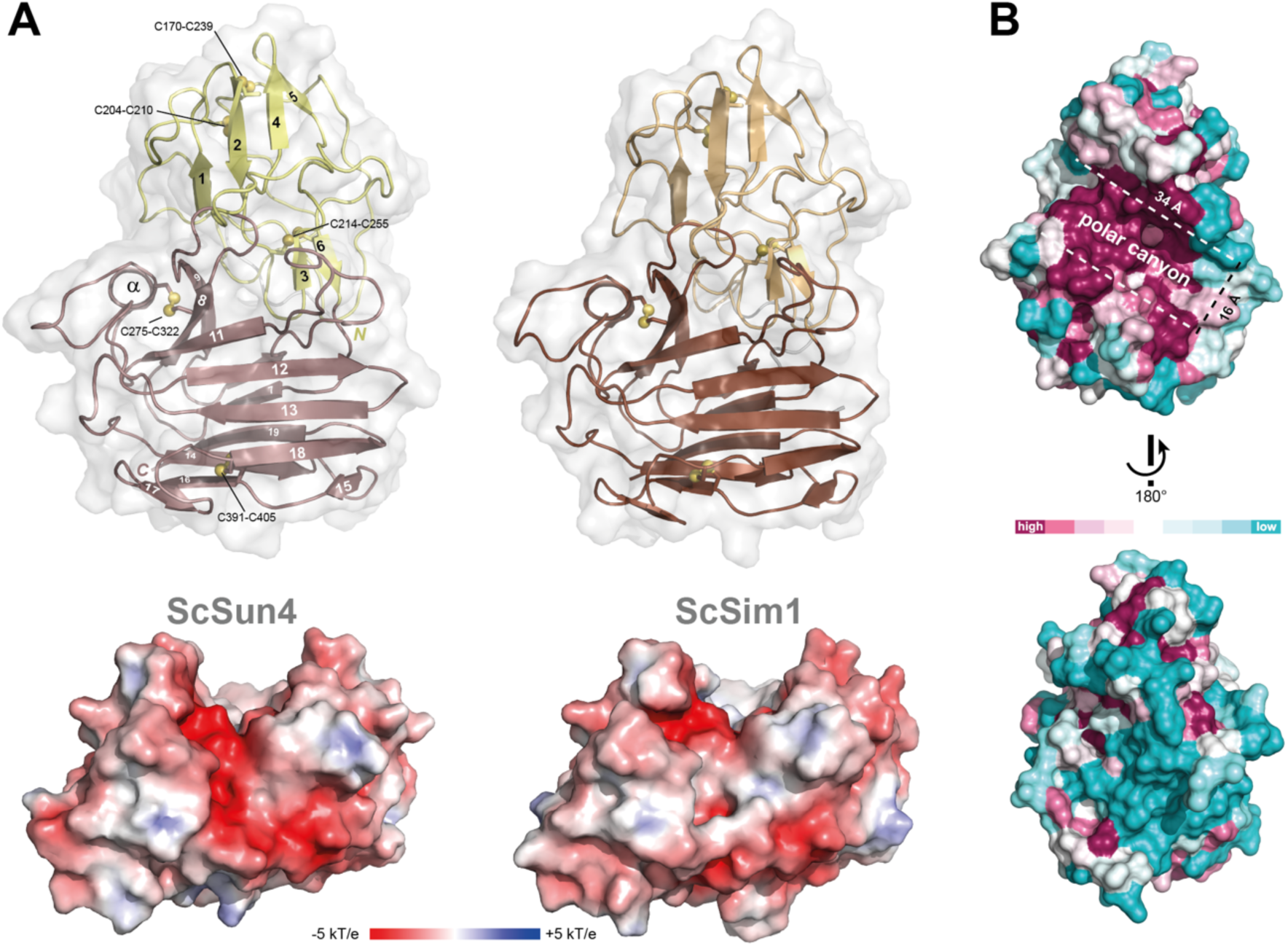
Structural features of the SUN domains from ScSun4 and ScSim1. A) Crystal structures of the SUN domains from ScSun4 (PDB: 9T47; UniProt: P53616) and ScSim1 (PDB: 9T4Q; UniProt:P40472) are shown in the upper part as cartoons and reveal a two-domain architecture consisting of an N-terminal sushi-like fold (upper part) and a thaumatin-like C-terminal fold (lower part). For ScSun4, the secondary structure elements, the ten cysteines forming five disulfide bonds (C170-C239, C204-C210, C214-C255, C275-C322, C391-C405) and the termini are indicated. Surface charge predictions for ScSun4 and ScSim1 are shown at the bottom. B) Identification of a highly conserved, canyon-shaped surface structure in fungal SUN domains. Evolutionary conservation of amino acid positions in the SUN domain of ScSun4 was determined by using the ConSurf Server (38) and 100 homologs collected from UNIREF90 (36) via CSI-BLAST (37) (E-value 0.0001). The maximal ID between the sequences used was set to 90 %, and to 35 % for homologs, respectively. The color-coded protein surface shows the predicted degree of conservation on the surface of ScSun4 ranging from high (maroon) to low (turquoise).

Our structure reveals a bilobal protein architecture consisting of two structural subdomains connected via a small linker unit with an overall size of 62.9 x 99.2 x 102.2 Å^3^ (PDB entry 9T47). The 11 kDa N-terminal subdomain is formed by a total of six β-strands and contains six cysteine residues that form three disulfide bonds (Figure 2A left, Figure S3A). Structure similarity search of this subdomain with foldseek (35) reveals a high similarity to the sushi domain (INTERPRO family IPR000436), a fold that is otherwise only found in metazoa. For example, the sushi fold of the interleukin 15 receptor α-subunit (IL-15Rα, PDB entry 2PSM) adopts the same topology despite only 9% sequence identity and a root mean square deviation (RMSD) of 2.57 Å for 54 C_α_-atoms (Figure S3B). The second subdomain of ScSun4 (V259-N420) consists of a total of 13 β-strands and two disulfide bonds (Figures 2A, S3A) and corresponds to the thaumatin-like fold (INTERPRO family IPR001938). For example, the thaumatin-like xylanase inhibitor TLXI (PDB entry 3G7M) matches the SUN4 thaumatin-like domain with a RMSD of 2.64 Å for 120 C_α_-atoms and a sequence identity of 15% (Figure S3B). Furthermore, the SUN domain of ScSun4 lacks the highly conserved REDDD-motif commonly found in thaumatin-like proteins (TLPs) .Notably, the combination of a sushi-like and a thaumatin-like fold uncovered by the crystal structure of the ScSun4 SUN domain represents a unique and previously unknown feature of fungal SUN family proteins.

We also solved the crystal structure of the SUN domain of ScSim1 (Figure 2A right, PDB entry 9T4Q), a closely related paralog of ScSun4 (85 % identity; Figure S1), at a resolution of 1.2 Å (Table S2). As expected, the structures of the ScSun4 and ScSim1 SUN domains resemble each other including the arrangement of the sushi- and thaumatin-like subdomains (RMSD 0.394 Å for 220 C_α_-atoms) and their disulfide bonds (Figure 2A). Furthermore, both SUN paralogs share highly acidic surfaces (Figure 2A, bottom), which is reflected by their calculated pI values of 4.1 (ScSun4) and 4.4 (ScSim1), respectively. Further analysis of the 77 SUN domains selected for the phylogenetic ML-tree analysis shown above (Figure 1B) reveals an average pI value of roughly 5.0, indicating a conservation of acidic surface properties throughout the phylum of Ascomycota. Nevertheless, the average pI values of Sun domains found within the subphyla of Saccharomycotina (n = 30; pI = 4.51) and Pezizomycotina (n = 42; pI = 5.24) significantly differ from each other, whereas their overall structures are highly similar (Figure S4).

Based on the crystal structures of ScSun4 and ScSim1, we next determined highly conserved surface residues within the fungal SUN domain family. For this purpose, we chose 100 homologous SUN domains from the UNIREF90 database (36) using the CSI-BLAST tool (37) for calculating a surface map by the ConSurf Server (38). This analysis reveals a highly conserved canyon-shaped surface between the sushi and the thaumatin subdomains (Figures 2B). This canyon has a length of ∼ 34 Å and a width of ∼ 16 Å, respectively, and contains a notable number of polar and aromatic surface residues. The dimension and surface properties of the highly conserved canyon are in line with the previously proposed idea that fungal SUN domains have a conserved β-glucan binding function that involves surface cavities with polar and aromatic properties (39).

### Identification of ScSun4 surface residues essential for *in vivo* function

To further analyze the function of the highly conserved polar canyon of fungal SUN domains, we performed a structure-based mutational analysis of ScSun4, because its absence in *S. cerevisiae* causes a specific cell clustering phenotype due to incomplete cell separation (24). However, because no quantitative assay for robust scoring the cell clustering phenotype was available, we first established a novel assay, named <u>Q</u>uantitative Cell <u>C</u>luster <u>A</u>nalysis (QCA), which allows to reliably determine the mean size of thousands of cell clusters formed by different yeast strains (Figure S5). As shown in Figure 3A, absence of ScSun4 causes pronounced cell clustering, a phenotype which can be reliably scored by the QCA assay with high statistical significance (Figure 3B and Table S3). Importantly, expression of the *ScSUN4* gene from a low-copy plasmid in a *sun4Δ* mutant strain is sufficient to complement the cell separation phenotype (Figure 3B and Table S3), which allowed to further analyze the effects of mutations in the SUN domain of ScSun4. Based on these results, we determined the *in vivo* function of a total of twelve conserved residues within the polar canyon of ScSun4. For this purpose, we individually exchanged five residues (T277, D278, Y279, E283, Q318) located at the bottom of the canyon for alanines, as well as seven residues (D230, R232, W310, W334, N343, W344, N367) on its side loops (Figures 3C and S1A). Plasmid-based expression of these mutants in a strain lacking the endogenous *ScSUN4* gene revealed, that the five variants carrying mutations at the bottom of the canyon fail to fully complement the cell clustering phenotype. This indicates that corresponding residues significantly contribute to the cell separation function of ScSun4. In contrast, individual mutations of either of the seven residues on the side of the canyon had no significant effect, because corresponding mutants are able to fully complement the cell clustering phenotype (Figure 3D and Table S4). However, combined mutation of either D230, R232, W310 (DRW) or W334, N343, W344, N367 (WNWN), respectively, significantly reduced cell separation, indicating that residues localized on the two sides of the canyon also contribute to the function of ScSun4.

**Figure 3.**
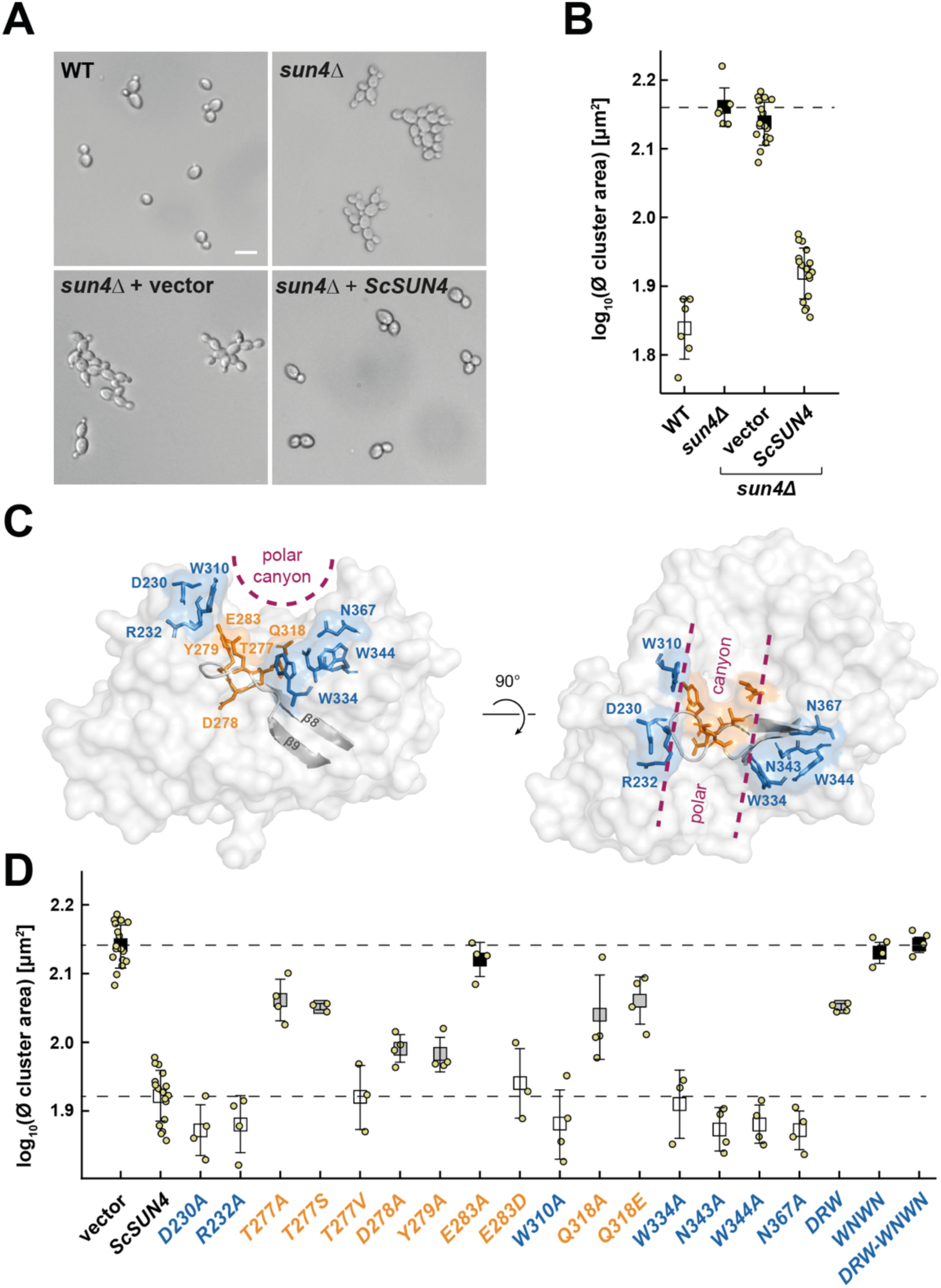
Structure-based mutational analysis of ScSun4. A) Microscopy. Effect of a ScSun4 deletion mutation on cell clustering was analyzed by bright-field microscopy of a yeast strain (YHUM3154) carrying the chromosomal *ScSUN4* gene (WT) and a yeast strain with a chromosomal *sun4Δ* mutation (YHUM3160), that was carrying either no plasmid (*sun4Δ*), or a control plasmid (vector pRS316) without *ScSUN4* (*sun4Δ* + vector), or a plasmid (BHUM3438) with *ScSUN4* (*sun4Δ* + *ScSUN4*), respectively. Before microscopy, strains were grown into logarithmic growth phase. White bar corresponds to 10 µm. B) Quantitative cell clustering analysis (QCA) of strains described in *A)*. Small squares indicate the average log_10_ values of at least three independent measurements of the mean cluster areas (yellow dots) for each strain, with error bars indicating the standard deviation. Statistical significance of the differences measured in comparison to the *sun4Δ* strain carrying no plasmid (dotted line) is shown as color-code according to the *P* values obtained by an unpaired t test (Table S3), with black squares indicating strains exhibiting *P* values > 0.01 and with white squares for strains with *P* values < 0.001, respectively. C) Structure of the polar canyon of ScSun4. The location of the canyon at the protein surface is indicated by dotted lines (maroon). Conserved residues localized at the bottom of the canyon (T277, D278, Y279, E283, Q318) are shown in orange and residues on its side loops (D230, R232, W310, W334, N343, W344, N367) are colored in blue. D) Effects on cell clustering by mutations of residues in the polar canyon. Yeast strains with a chromosomal *sun4Δ* mutation and carrying either a control plasmid (vector) or a plasmid with *ScSUN4* (*SUN4*) or a plasmid with the indicated *ScSUN4* mutational variant, respectively, were quantified for cell clustering by QCA. Statistical significance of differences measured in comparison to the *sun4Δ* strain carrying the non-mutated *ScSUN4* variant (dotted line) is color-coded according to the *P* values obtained by an unpaired t test (Table S4). Black and grey squares are shown for strains exhibiting a mean cluster area above that of the *ScSUN4* control strain and with *P* values < 0.001 (black), or *P* values between 0.01 and 0.001 (grey), respectively. White squares are displayed for strains exhibiting a mean cluster area above that of the *ScSUN4* strain and with *P* values > 0.01, or for strains with a mean cluster area below that of the control. *ScSUN4^DRW^* corresponds to ScSUN4*^D230 R232A W310A^*, ScSUN4*^WNWN^* to ScSUN4*^W334A N343A W344A N367A^*, and ScSUN4*^DRW-WNWN^* to ScSUN4^D230A R232A W310A W334A N343A W344A N367A^, respectively.

To further analyze the importance of specific amino acid side chain features, we characterized a number of mutations at the position of the three residues (T277, E283, Q318), whose mutations cause the strongest cell separation defects (Figure 3D) without affecting the overall ScSUN4 structure as shown for its E283A and Q318A mutants (PDB entries: 9T4O, 9T4N). This analysis shows that expression of ScSun4^T277V^, but not ScSun4^T277S^, complements the cell separation phenotype. This indicates that the methyl group of T277 is important for functionality, but not its hydroxyl group. We furtherthat an acidic residue at position 283 substantially contributes to proper function, because expression of ScSun4^E283D^ partially restores cell separation. Finally, mutation of the glutamine at position 318 to glutamate leads to a non-functional variant, indicating that the amide group of Q318 is required.

In summary, our mutational analysis defines the highly conserved polar canyon of ScSun4 as a central functional region of its SUN domain.

### Functional analysis of the SUN domain family of *S. cerevisiae*

Having established the structural basis for the ScSun4 function, we next sought to determine the cell separation function of the SUN domains of all other SUN family members of *S. cerevisiae*, representing the ascomycetal subphylum of Saccharomycotina (budding yeasts). Although previous studies have shown an involvement of ScSun4 and ScUth1 in this process (24), no quantitative analysis has been performed comparing all five family members side by side. We therefore constructed a set of isogenic *S. cerevisiae* mutant strains lacking either of the *ScSUN4*, *ScSIM1*, *ScUTH1* or *ScNCA3* genes, which encode group I SUN domains, or the group II gene YMR244w (www.yeastgenome.org), respectively, that we named *ScADG3* in order to emphasize its phylogenetic relationship to other fungal group II members (Figures 1 and S1). Comparative analysis of the cell separation phenotype of these strains by QCA revealed that unlike *ScSUN4* no significant cell clustering is induced by the absence of any other SUN family gene (Figure 4, Table S5). Thus, ScSun4 appears to play the dominant role in cell separation during cell division in *S. cerevisiae*.

**Figure 4.**
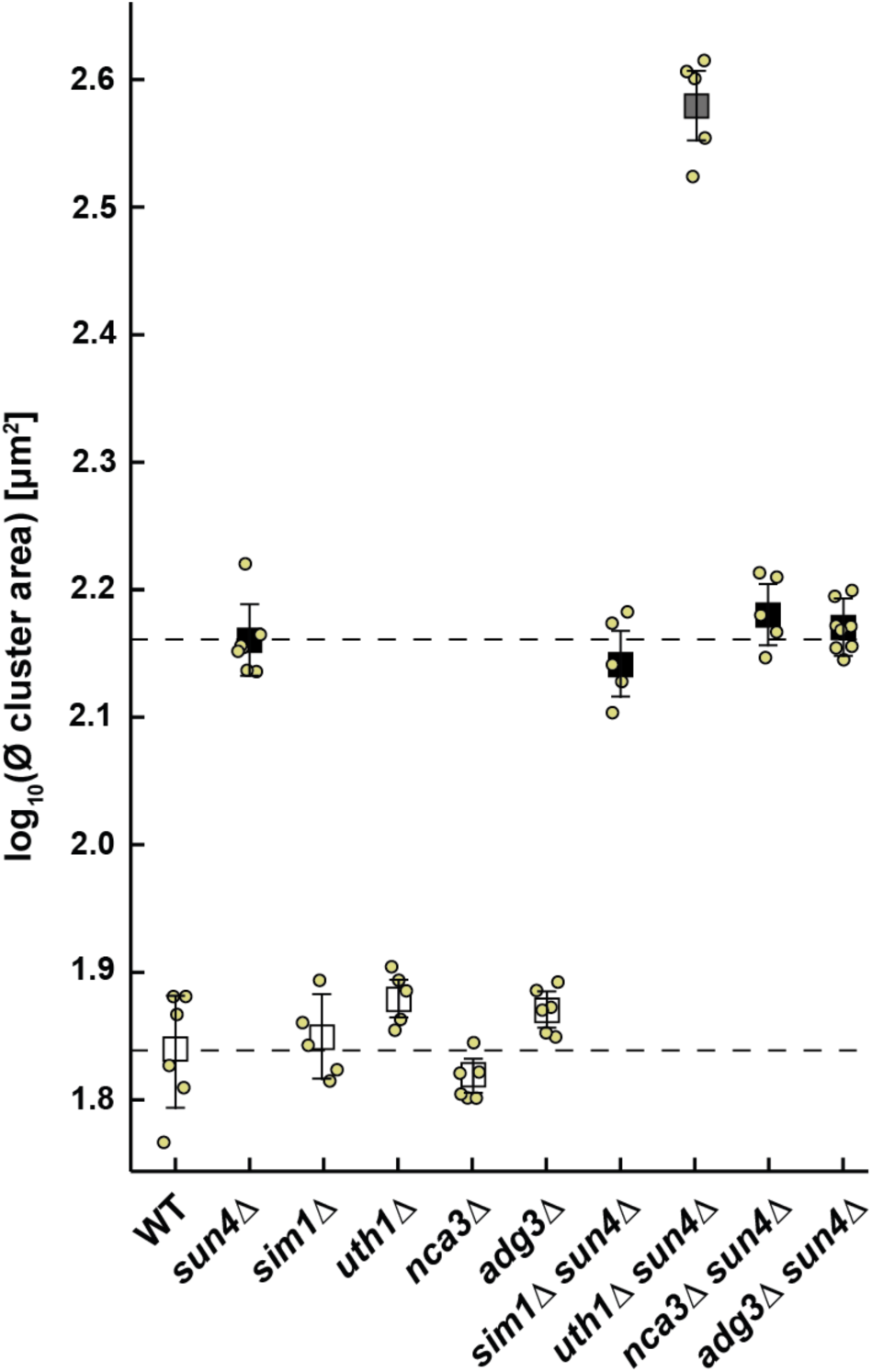
Functional analysis of the SUN domain family of *S. cerevisiae*. Isogenic yeast strains carrying the indicated chromosomal deletions of *ScSUN4* (*sun4Δ*), *ScSIM1* (*sim1Δ*), *ScUTH1* (*uth1Δ*), *ScNCA3* (*nca3Δ*) or *ScADG3* (*adg3Δ*), respectively, were quantified for cell clustering by QCA. A a control, an isogenic wild-type (WT) strain is shown. Small squares indicate the average log_10_ values of at least three independent measurements of the mean cluster areas (yellow dots) for each strain, with error bars indicating the standard deviation. Statistical significance of the differences measured is color-coded according to the *P* values obtained by an unpaired t test (Table S5) and in comparison to the WT strain and/or the *sun4Δ* strain (dotted lines). White squares are shown for *P* values > 0.02 in comparison to the WT strain. Black squares indicate *P* values > 0.02 in comparison to the *sun4Δ* strain. Grey squares are shown for *P* values < 0.001 in comparison to both the WT and the *sun4Δ* strain.

In order to uncover possible minor roles of the other family members, we further constructed a set of double mutant strains that lack *ScSUN4* in combination with any of the other SUN genes. Phenotypic analysis of these strains revealed that absence of both, *ScSUN4* and *ScUTH1*, significantly enhances cell clustering when compared to the phenotype observed for the *sun4Δ* single mutant (Figure 4, Table S5). In contrast, no such additive effects were observed for the other double mutant strains. This indicates that ScUth1 is able to confer cell separation functionality, a finding that is supported by previous observations (24).

The fact that all SUN family proteins of *S. cerevisiae* consist of a conserved SUN domain fused to a non-conserved C-terminal domain, prompted us to test to what extent SUN domains would be functionally exchangeable between different members with respect to cell separation. For this purpose, we constructed four variants of the *ScSUN4* gene that carry the SUN domain of either *ScSIM1*, *ScUTH1*, *ScNCA3* or *ScADG3*, respectively, instead that of *ScSUN4* (Figures 5A and S1A). The four chimeric variants were then expressed in the *S. cerevisiae sun4Δ* strain, to test the ability of these constructs to restore the specific cell separation defect of this mutant. By using the QCA assay, we found that expression of the two chimeric variants carrying the SUN domain of either *ScSIM1* (group I), which shares 85 % identity with that of ScSun4, or that of *ScUTH1* (62 % identity; group I), were able to almost fully complement the absence of the endogenous *ScSUN4* gene (Figure 5B and Table S6). In contrast, the SUN domains of either *ScNCA3* (61 % identity; group I) or *ScADG3* (32 % identity; group II) confer only very weak cell separation functionality in the context of *ScSUN4*. These data show that group I SUN domains are in principle functionally interchangeable between *S. cerevisiae* paralogs, indicating a conserved molecular function of certain, but not all SUN domains. Also, functional replacement does not seem to merely depend on phylogenetic distances.

**Figure 5.**
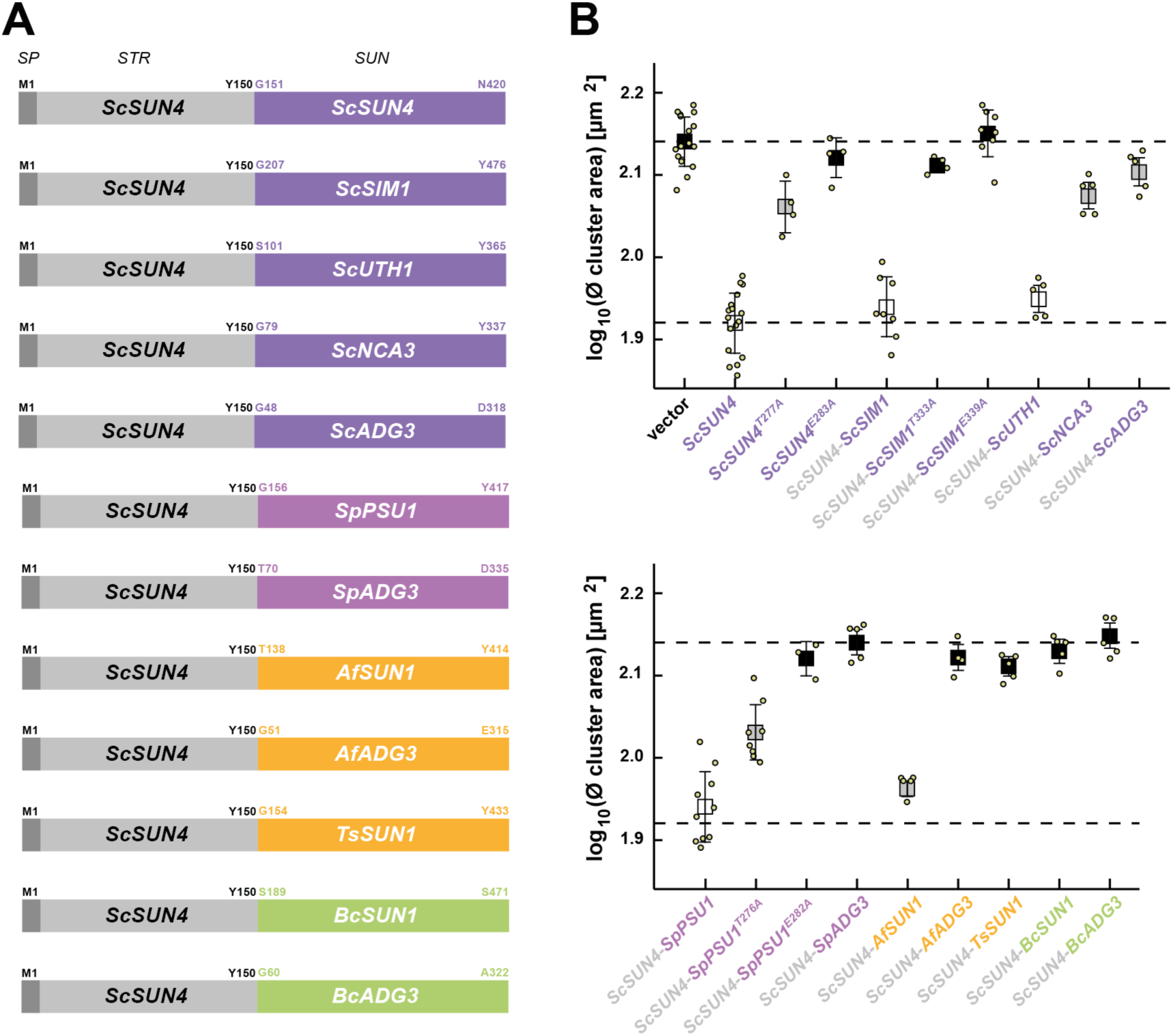
Comparative functional analysis of SUN domains from budding yeast, fission yeast and filamentous fungi. A) Domain structure of *ScSUN4* and chimeric variants. The endogenous *ScSUN4* gene consists of a signal peptide for secretion (SP) and a serine-threonine-rich region (STR), which together encompass the N-terminal part (M1-Y150), followed by a C-terminal SUN domain (G151-N420). The diverse chimeric variants consist of the N-terminal part of *ScSUN4* fused to the SUN domain of either *ScSIM1* (G207-Y476), *ScUTH1* (S101-Y365), *ScNCA3* (G79-Y337), *ScADG3* (G48-D318), *SpPSU1* (G156-Y417), *SpADG3* (T70-D335), *AfSUN1* (T138-Y414), *AfADG3* (G51-E315), *TsSUN1* (G154-Y433), *BcSUN1* (S189-S471) or *BcADG3* (G60-A322). *Sc*: *S. cerevisiae*; *Sp*: *S. pombe*; *Af*: *A. fumigatus*; *Ts*: *T. stipitatus*; *Bc*: *B. cinerea*. B) Quantitative cell clustering analysis of *sun4Δ* strains expressing plasmid-borne SUN domain variants shown in (*A*) or versions carrying the indicated point mutations. As controls, values for strains carrying either an empty vector or the non-mutated *ScSUN4* variant are shown and indicated by dotted lines. The average log_10_ values of at least three independent measurements of the mean cluster areas (yellow dots) for each strain are shown as small squares, with error bars indicating the standard deviation. Statistical significance of the differences measured is color-coded according to the *P* values obtained by an unpaired t test (Table S6) and in comparison to the *sun4Δ* strain carrying the non-mutated *ScSUN4* variant. Black and grey squares are shown for strains exhibiting a mean cluster area above that of the *ScSUN4* control strain and with *P* values < 0.001 (black), or *P* values between 0.01 and 0.001 (grey), respectively. White squares are displayed for strains exhibiting a mean cluster area with *P* values > 0.01.

To further refine the above conclusion, we determined the spatial topology of the five conserved residues at the bottom of the polar canyon of the ScSim1 SUN domain (Figure S1A), which confer functionality in ScSun4 (Figure 3D and Table S4). Comparison of the crystal structures of ScSun4 and ScSim1 reveals that the topologies of these residues are highly similar, indicating a conserved function. To further test this assumption, we constructed two additional chimeric variants in the *ScSUN4* context, *ScSIM1*^T333A^ and *ScSIM1*^E339A^, that carry alanine mutations at the positions corresponding to T277 and E283 in ScSun4 (Figure S1A). As shown in Figure 5B, both of these mutations in the ScSim1 SUN domain cause cell clustering to a degree, which is comparable to that caused by the respective mutations in ScSun4, indicating a conserved function of residues at the bottom of the polar canyon.

### Functional analysis of SUN domains from fission yeast and filamentous fungi

To investigate the function of SUN domain proteins of Taphrinomycotina (fission yeasts) and Pezizomycotina (filamentous fungi), we performed a functional analysis of seven SUN domains from four species of these ascomycetal subphyla. Specifically, we constructed seven further chimeras of the *ScSUN4* gene, which carry the SUN domain of either *S. pombe SpPSU1* (53 % identity to ScSUN4; group I) or *SpADG3* (31 %/group II), *A. fumigatus AfSUN1* (43 %/group I) or *AfADG3* (32 %/group II), *B. cinerea BcSUN1* (39 %/group I) or *BcADG3* (31 %/group II), or *T. stipitatus TsSUN1* (40 %/group I), respectively (Figures 5A and S1A). Again, these chimeric variants were functionally analyzed by the QCA assay upon expression in a *S. cerevisiae sun4Δ* strain. Here, we found that the group I SUN domains from *SpPSU1* and *AfSUN1* were able to functionally replace the respective *ScSUN4* domain, whereas the other SUN domains tested were found to confer no significant function (Figure 5B and Table S6). Thus, certain group I SUN domains from the subphyla of both Taphrinomycotina (*SpPSU1*) and Pezizomycotina (*AfSUN1*) are able to confer *S. cerevisiae* cell separation in the context of ScSUN4. This finding further supports the view, that group I SUN domains with significant phylogenetical distance still possess a conserved molecular function. In contrast, none of the tested group II domains were found to complement the function of the group I domain from ScSun4.

We corroborated this conclusion by determining the spatial topology of conserved residues at the bottom of the polar canyons of SpPsu1 and AfSun1. Alphafold3 models of the SUN domains of SpPsu1 and AfSun1 revealed that their overall structure as well as the topology of conserved residues at the bottom of their polar canyon are highly conserved. Analogously to the *ScSIM4* variants, we constructed two additional *ScSUN4* chimeras harboring either the *SpPSU1*^T276A^ or *SpPSU1*^E282A^ domain (Figure S1A). As found for *ScSUN4* and the *ScSIM1* chimera, both mutations in the *SpPSU1* SUN domain significantly abrogated its cell separation function in *S. cerevisiae* (Figure 5B).

Taken together, these results support the conclusion that the polar canyons of phylogenetically very distant group I SUN domains appear to possess a conserved biochemical function that confers cell separation.

### Structural evidence for binding of β-1,3-glucan polymers by group I SUN domains

The finding of a broadly conserved function of diverse group I SUN domains in cell separation prompted us to analyze their interaction with β-glucans. SPR spectroscopy of ScSun4 with laminarin, a soluble β-1,3-glucan, shows only weak affinity (K_D_ = 3.6 mM, Figure S6). However, one may note that laminarin differs from the fungal β-1,3-glucans by interspersed β-1,6-sections. Furthermore, much of the β-1,3-glucans of fungal cell walls consist of single-stranded and triple-helical structures (12), with the latter being stabilized by hydrogen-bonding. Alphafold3 models of the ScSun4 domain and single-stranded β-1,3-glucans show binding poses, where the glucan fills the polar canyon spanned between the sushi- and thaumatin-like domains (Figure 6A). These binding poses are also reproducible for other members of the group I SUN domains including ScSim1 (Figure 6A, bottom). In addition to the floor region of the polar canyon, a significant fraction of interactions appears to be contributed by four adjacent loop regions (β_3_-β_4_, β_8_-β_9_, β_11_-β_12_, β_13_-β_14_). The dimension of the canyon is also compatible for accommodating triple-helical β-1,3-glucans, such as insoluble curdlan. Accordingly, computational docking places the right-handed triple helix of curdlan well into the polar canyon (Figure 6B), with an analogous contribution by the aforementioned loop regions. However,docking is indecisive with regard to the orientation of the trimeric glucan. Nevertheless, molecular dynamics analysis shows the stability of the docked complexes within the simulation periods of 100 ns. The volumetric density map of the bound triple glucan showing the minimal and maximal probability density reveals no major dislocations of the bound ligand apart from occasional kinking events leading to a temporal release from the β_3_-β_4_ loop of the sushi-like fold (Figure 6C).

**Figure 6.**
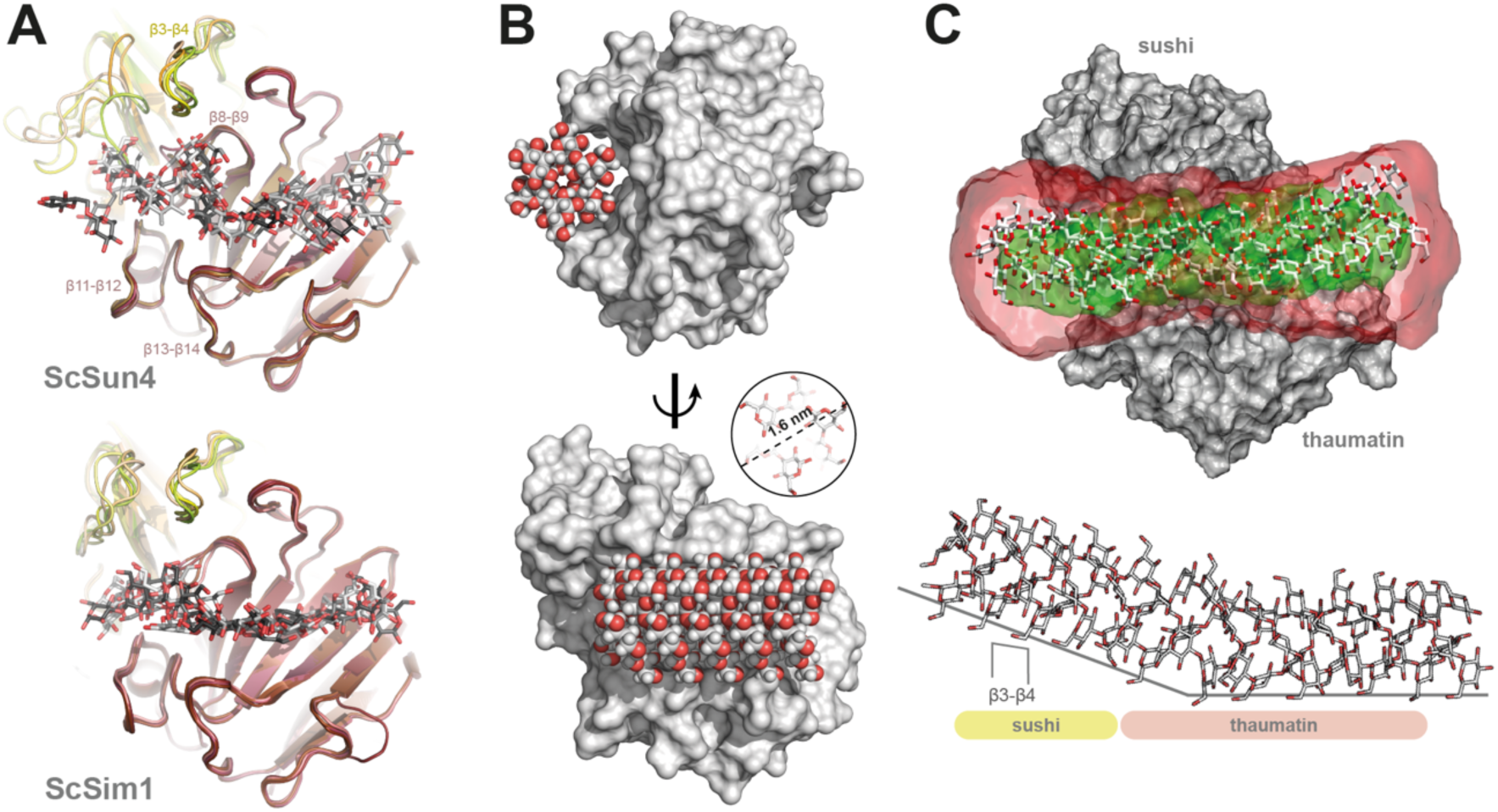
Binding of β-1,3-glucans to the polar canyon of ScSun4. A) Models of ScSun4 (top) complexed to laminarioctaose as predicted by Chai-1 (96). The four loops in direct interaction with the β-1,3-glucan are indicated. For comparison the predicted complexation of these linear β-1,3-glucans with ScSim1 in the carbohydrate-binding polar canyon is shown below. B) Triple-helical β-1,3-glucan (Curdlan, DP20) is found by *in silico* docking within the ScSun4 polar canyon (ScSun4 surface: grey, glycans: CPK models) as well. The inlet shows the diameter of the triple-helical β-glucan (1.6 nm). C) VMD-based volumetric density map calculated with the VolMap Plugin 1.1 and the structures of triple-helical β-glucan and the ScSun4 SUN domain (upper part). All frames (100 ns) were computed and combined using minimum (green) and maximum (red) occupancy showing a dumbbell-shaped map. In later simulation periods, kinking of triple-helical β-glucan occurred, but without the loss of the ligand (lower part). Kinking is induced by steric repulsion of triple-helical β-glucan by the sushi loop β_3_-β_4_. Sushi-like and thaumatin-like subdomains of ScSun4 are indicated.

Taken together, these data show that the structures of group I SUN domains are well-suited to accomodate both single-stranded as well as triple-helical β-glucan structures.

## Discussion

In this study, we find that fungal SUN domains of both group I and group II consist of fusions between a thaumatin-like and a sushi-like fold, which together define a highly conserved polar canyon, that is well fitted to bind mono-as well as triple-helical β-glucans. We further find that group I SUN domains of diverse fungi are able to confer cell separation, a function that crucially depends on specific structural features of the polar canyon and that cannot be fulfilled by group II SUN domains. As such, our study provides novel and detailed evidence supporting the view that fungal SUN domain proteins are involved in reorganization of the β-glucan network, specifically with respect to the separation of individual cells.

A central finding of our structural analysis is that the core function of fungal SUN domains is enabled by combining a thaumatin-like with a sushi-like fold. Several thaumatin-like proteins (TLP) have previously been described in eukaryotes such as animals, plants and fungi, where TLP are associated with either glycan binding particularly of β-1,3-glucans, enzyme inhibition of enzymes such as xylanases, alpha-amylase or trypsin, pathogen resistance or membrane permeabilization (40–45). Other functional properties have been reported for TLP, including binding to proteins such as actin, viral CMV-1 protein, yeast glycoproteins and G-Protein Coupled Receptor (GPCR) or to hormones such as cytokinins (46). Most typical TLP described to date are small proteins with a typical weight ranging from 20-26 kDa harboring 16 conserved cysteine residues that form eight disulfide bonds (47). Numerous TLP structures have been deposited in the protein data bank yet, most of which are showing a strongly conserved structural organization with a characteristic polar canyon domain that comprises the highly conserved REDDD-motif (48). Besides that, small TLP (sTLP) have been identified in conifers and monocots. These show only ten conserved cysteine residues that form five disulfide bonds and despite the good conservation of the amino acids in primary sequence in sTLP, but they do not organize into a polar canyon that was shown for the non-competitive xylanase inhibitor TLXI (49). With respect to sushi-like domains, previous studies have revealed that most known sushi/SCR/CCP domains show a stalk-like structure and have been assigned a substrate recognition function (50). In addition, sushi-like domains have been proposed to function as rigid spacers or handles that help to position other domains for activation or exertion of their catalytic or binding function, as exemplified for the chymotrypsin-like serine protease (SP) (51). Our finding that sushi-like domains are fused to thaumatin-like domains in all known SUN family proteins therefore suggests, that the biochemical function of the thaumatin-like moiety is supported or might be regulated by the sushi-like part.

A further outcome of our study is that the polar canyon of fungal SUN domains is well fitted to bind unbranched single as well as triple-helical β-1,3-glucans. So far, a small number of proteins from animals and bacteria have been demonstrated to show three-dimensional structural characteristics for the binding of such polymers (39, 52). In insects, the β-glucan recognition protein (βGRP) is a crucial pattern-recognition receptor that specifically binds β-1,3-glucans of fungal cell walls (53).

The βGRP consists of an N-terminal β-1,3-glucan-recognition domain and a C-terminal glucanase-like domain, with the former having a β-sandwich fold, often seen in carbohydrate-binding modules. Both NMR titration experiments as well as crystallographic structures of βGRP-N indicate that this domain specifically recognizes three structured laminarihexaoses mimicking a branched triple-helical structure of β-1,3-glucan (54, 55). Two further structurally well-studied bacterial multidomain proteins, which were implicated in triplex β-glucan-binding, include a family 81 glycosyl hydrolase (GH81) β-1,3-glucananase (BH0236) from *Bacillus halodurans* (56) and a GH64 family β-1,3-glucanase (PbBgl64A) from *Paenibacillus barengoltzii* (57). BH0236 is composed of three parts: an N-terminal GH81 catalytic module, an internal family 6 carbohydrate binding module (CBM6) that binds to the non-reducing end of β-1,3-glucan chains, and a C-terminal CBM56 binding to β-1,3-glucan chains (56). In contrast, PbBgl64A is composed of two regions: a N-terminal CBM56 with the ability to bind β-1,3-glucans and a C-terminal region that corresponds to a GH64 β-1,3-glucanase domain, which belongs to the GH64-TLP superfamily including both GH64 as well as thaumatin-like proteins (57). Despite these differences in primary structure, crystal structures of the two bacterial enzymes bound to diverse long and short β-1,3-glucans indicate that the architecture of the catalytic sites of both of these proteins are prone to accommodate the triple helical quaternary structures of β-1,3-glucan chains. Our present structural analysis of the fungal-specific SUN protein family reveals the existence of a further, previously uncharacterized domain architecture, which appears to be prone to recognize both single as well as triple helical β-1,3-glucan chains. As such, our study describes a novel structural solution for this biochemical task, which separately evolved in the domain of fungi. The precise molecular function of SUN domain proteins, however, is not completely clear. A previous study reported a weak glucanase activity for AfSun1 assigned as GH132 family (26). Nevertheless, we were unable to assign an active glucanase site within the polar canyon, leaving other possibilities for SUN domain functions like acting as adapters between β-1,3-glucans and other cell wall proteins.

With respect to the evolution and diversification of SUN domains, it was previously recognized that in comparison to group I SUN domains, most group II family members possess an additional C-terminal serine/threonine (S/T)-rich domain (24–26). Our current study reveals further significant structural and functional distinctions between group I and group II. Sequence comparison using EFI-EST demonstrates a clear separation between group I and group II SUN domains. In addition, our functional assays indicate that the function of the group I SUN domain of ScSun4 cannot be substituted by any tested group II domain. Moreover, our phylogenetic analysis reveals that all group I domains known so far are found in Ascomycota, whereas the few SUN domains found outside this phylum, in Mucoromycota or Zoopagomycota, belong to group II (Figure S2). These findings, together with the fact that most ascomycetes contain SUN domains of both groups, support the view that (i) the ancestor of the SUN domain family was similar to group II and that (ii) a diversification into group I and group II occurred early within the phylum of Ascomycota, which included the loss of the C-terminal serine/threonine (S/T)-rich domain.

Our analysis also highlights class-specific distribution patterns of group I and group II SUN domains across different ascomycetal subphyla. In filamentous fungi of the subphylum Pezizomycotina, the number of SUN domain family members is typically restricted to one or two. Most species in this group possess either a single member, predominantly from group I, or two members, often representing both group I and group II. These patterns might indicate an evolutionary constraint specific to filamentous lifestyles or an adaptation to certain ecological niches. Organisms with a single member may for instance rely on a streamlined SUN domain function, possibly linked to cell wall integrity or basic morphogenesis, which are critical for filamentous growth. In contrast, the presence of both group I and group II domains could indicate a division of labor, with one group potentially contributing to more specialized processes such as alternate lifestyles or adaptation to specific environments. Nevertheless, class-specific distribution patterns do not necessarily reflect the precise biochemical function of group I or group II domains. For example, our functional analysis has shown that the group I domain of AfSun1 from the class of Eurotiomycetes is able to complement the function of ScSun4, whereas the group I domain of TsSun1, belonging to the same taxonomic class, cannot. Together with the fact that these two group I SUN domains share only 54 % sequence identity (Figure S1), this finding demonstrates further functional diversification of SUN domains within a given class. A possible explanation might be a structural adaptation of SUN domains to different types of β-glucans, which can vary in filamentous fungi (6, 9, 13, 39, 58, 59). In contrast to Pezizomycotina, yeast-like organisms from the subphyla Saccharomycotina and Taphrinomycotina tend to exhibit an increased number of group I family members, with some organisms carrying three or more paralogs (Figure S2C). This disparity implies that group I SUN domains may have undergone expansion and functional diversification in these yeast-like fungi, while group II paralogs remained relatively stable in number. For Saccharomycotina and Taphrinomycotina, the expansion of group I members might reflect an evolutionary strategy to diversify functions in rapidly changing environments, such as fermentation or pathogenicity, where multiple paralogs could fine-tune responses to varying substrates or conditions. The limited presence of group II genes in these subphyla also suggests that they may serve a conserved, essential role less prone to duplication. An increased number of SUN domain genes might further be required for the yeast-like growth pattern, because the budding yeast lifestyle is well-known to consist of several distinct growth phases that include profound cell wall rearrangements such as (i) initial polar cell growth after bud emergence, (ii) a switch from polar to isotropic growth at the G2/M boundary of the cell division cycle, and (iii) a redirection of cell wall component secretion towards the neck region, to permit the degradation of septum carbohydrates between dividing cells leading to cell separation (60). Here, diverse SUN domain members might have evolved for an optimized temporal or spatial remodeling of the β-glucan network during yeast-like cell growth.

With respect to functional diversification of SUN domain paralogs in yeast-like fungi, our study provides novel insights at the example of the budding yeast *S. cerevisiae*, which harbors five family members: ScSun4, ScSim1, ScUth1, ScNca3 and ScAdg3. So far, genetic analysis has uncovered an involvement of ScSun4 and ScUth1 in cell wall biogenesis and septation (24, 28), whereas no specific cell wall phenotypes have been described for yeast mutants lacking either ScSim1, ScNca3 or ScAdg3 (www.yeastgenome.org). In addition, several genetic studies have implicated ScUth1, ScSun4 and ScSim1 in mitochondrial biogenesis (61–63), as well as resistance to toxins (64, 65) and other stressors including ethanol (66, 67) or diverse cell wall-perturbing agents (28, 65, 68). However, many of the effects observed in mutants lacking these SUN family proteins on mitochondrial function and yeast physiology appear to be indirect and might be better explained by their role in cell wall biogenesis (28). This view is underscored by the fact that ScSun4, ScSim1 and ScUth1 are secreted proteins that are found in the extracellular space (68). Our current study supports the view that ScSun4, ScSim1 and ScUth1 play important roles in cell wall reorganization, and our data obtained with ScSun4 chimeras demonstrates that they share a common biochemical function that is likely to affect the β-glucan network. Nevertheless, the cellular employment of ScSun4, ScSim1 and ScUth1 functions might well differ both temporally and spatially. This conclusion is supported by the diverse phenotypic patterns observed in mutants carrying individual gene deletions (as discussed above), and by the fact that according to the SPELL database (https://spell.yeastgenome.org), the expression patterns of the *ScSUN4*, *ScSIM1* and *ScUTH1* genes are similar, but not identical. Clearly, future detailed studies are required to uncover potential differences in temporal expression patterns encoded in the promoters of these genes. With respect to specific spatial patterns, a previous study has found Sun4 to be localized at the mother-daughter neck region (69). Our current study supports the view that the diverse N-terminal parts of SUN family proteins might confer specific cellular localization patters, given the fact that fusion of the N-terminal part of ScSun4 to the SUN domains of ScSim1 or ScUth1 create chimeras that fully complement the absence of ScSun4, whereas deletion of ScSim1 or ScUth1 alone does not cause a cell separation phenotype. Here, further detailed studies are required that investigate the precise function of the diverse N-terminal portions of these SUN domain proteins. With respect to the fourth *S. cerevisiae* type I SUN domain protein, ScNca3, our study reveals that its SUN domain is not able to complement the one of ScSun4 with respect to cell separation and thus might fulfill a biochemical function that differs from ScSun4, ScSim1 and ScUth1. This view is in agreement with the previous finding that in contrast to the other type I SUN domain proteins of *S. cerevisiae*, ScNca3 is not released into the extracellular space (68) and by the fact that the expression pattern found for *ScNCA3* significantly differs from the patterns of *ScSUN4*, *ScSIM1* and *ScUTH1* (https://spell.yeastgenome.org). Furthermore, ScNca3 possesses three unique charged surface residues (D176, E177, K181) in the β6/β7 region of its SUN domain, which are not found in the other type I SUN proteins of *S. cerevisiae* (Figure S1) and that might confer functional specificity. Finally, our study has uncovered that the single type II SUN domain protein ScAdg3 appears to fulfill a function in *S. cerevisiae* that differs from that of *ScSUN4*, *ScSIM1* and *ScUTH1*, a conclusion that is supported by the fact that expression pattern of *ScADG3* significantly differs from the patterns of the other *S. cerevisiae* SUN family members (https://spell.yeastgenome.org). As in the case of ScNca3, however, no specific phenotype has been described for mutant strains lacking *ScADG3* (www.yeastgenome.org). Clearly, future studies are needed to further characterize the specific functions of different SUN family proteins in *S. cerevisiae,* which might include more complex genetic approaches in combination with screening efforts for novel phenotypes. Such investigations might profit from more recent methods such as solid-state NMR of fungal cell walls (59, 70), in order to further elucidate how SUN domain proteins shape the cell wall architecture and offer deeper insights into their physiological roles and evolutionary diversification.

## Materials and Methods

### Yeast strains and growth conditions

All yeast strains used in this study are of the ∑1278b genetic background and listed in Table S7. Yeast strain YHUM3154 (*MAT***a**/*MATα*) was obtained by crossing of haploid yeast strains YHUM470 (*MAT***a**) and YHUM471 (*MATα*). Haploid yeast strains YHUM3107, YHUM3108, YHUM3101, YHUM3103, YHUM3104, YHUM3106, YHUM2881, YHUM2883, YHUM3605 and YHUM3607, were obtained by introduction of chromosomal deletions of *SUN4*, *SIM1*, *UTH1*, *NCA3* or *ADG3* in yeast strains YHUM470 and YHUM471, respectively, by using *sun4Δ::hphNT1*, *sim1Δ::kanMX6*, *uth1Δ::natNT2*, *nca3Δ::kanMX6* or *adg3Δ::kanMX6* deletion cassettes amplified by PCR from plasmids pFA6a-kanMX6, pFA6a-hphNT1 or pFA6a-natNT2 (71, 72). All genomic mutations were verified by Southern and/or PCR-based analysis. Diploid yeast strains YHUM3160, YHUM3156, YHUM3115, YHUM3602, YHUM3630, YHUM3162, YHUM3164, YHUM3642 and YHUM3637, carrying homozygous chromosomal deletions of *SUN4*, *SIM1*, *UTH1*, *NCA3* or *ADG3*, respectively, or combinations thereof, were obtained by crossing of appropriate haploid mutant strains. Standard methods for yeast culture medium and transformation were used as described (73). Plasmid carrying yeast strains used for *in vivo* functional analysis were obtained by fresh transformation of yeast strains of suitable chromosomal genotype with appropriate plasmids isolated from *E. coli*.

### Construction of plasmids

Plasmids used in this study are listed in Table S8. All plasmids constructed in this work were verified by DNA sequence analysis. Plasmids pET-28a(+)-ScSUN4 and BHUM3442 (pET-28a-*ScSIM1*) were obtained by PCR amplification of the SUN domains of the *ScSUN4* gene (encoding residues G147-N420) and of *ScSIM1* (encoding residues G202-N476) from *S. cerevisiae* genomic DNA using appropriate primers and insertion of the resulting DNA fragments into the pET-28a(+) Novagen expression vector (Merck, Darmstadt, Germany) using *Nde*I and *Xho*I restriction sites. Plasmids pET-28a-ScSUN4^E283A^ and pET-28a-ScSUN4^Q318A^ were obtained by site-directed mutagenesis of plasmid pET-28a(+)-*ScSUN4*). Plasmid BHUM3454 (pET-28a-SpPsu1) was constructed by PCR amplification of the SUN domain from the *SpPSU1* gene (encoding residues G165-Y417) from *S. pombe* genomic DNA and *Nde*I/*Xho*I-mediated insertion into pET-28a(+). Plasmid BHUM3438 was obtained by PCR amplification of a 2’932 bp DNA fragment carrying the *SUN4* gene from *S. cerevisiae* genomic DNA and insertion of the fragment into the centromere-based yeast vector pRS316 (74) using *Xba*I and *Bam*HI restriction sites, after *Eco*RV-mediated subcloning of the fragment in plasmid pJET1.2 (Fisher Scientific GmbH, Schwerte, Germany). Plasmids BHUM3657 to BHUM3659, BHUM3661 to BHUM3672 and BHUM3680, respectively, were obtained by site-directed mutagenesis of plasmid BHUM3438.

Plasmid BHUM3446 was generated by *in vivo* ligation in *S. cerevisiae*. For this purpose, a DNA fragment of the SUN domain (encoding residues G207-N476) of the *ScSIM1* gene was amplified from *S. cerevisiae* genomic DNA by PCR, thereby introducing flanking sequences with homology to the end of the STR region of ScSun4 (up to position Y150) and the terminator region of the *ScSUN4* gene, respectively. The resulting fragment was co-transformed together with the *Hpa*I-linearized plasmid BHUM3438 (pRS316-*ScSUN4*) into competent yeast cells to assemble a circular plasmid by homologous recombination. The resulting plasmid (BHUM3446) carrying the desired *ScSUN4-ScSIM1* chimeric gene was extracted from yeast and verified by DNA sequence analysis after cloning in *E. coli*. Plasmid BHUM3444 was generated by *in vivo* cloning following the same strategy used for plasmid BHUM3446, but using a DNA fragment of the SUN domain (encoding residues G156-N417) of the *SpPSU1* gene amplified from *S. pombe* genomic DNA. Plasmids BHUM3676 and BHUM 3677 were obtained by site-directed mutagenesis of plasmid BHUM3446, and plasmids BHUM3678 and BHUM 3679 were generated by site-directed mutagenesis of plasmid BHUM3444, respectively. Plasmids BHUM4093, BHUM4063, BHUM4043, BHUM4044, BHUM4042, BHUM4069, BHUM4046, BHUM4065 and BHUM4067 were generated by *in vivo* cloning in *S. cerevisiae* following the same strategy used for plasmid BHUM3446, but using synthetic and codon-optimized DNA fragments (Twist Bioscience, San Francisco, CA, USA) for the SUN domains of *ScUTH1* (S101-Y365), *ScNCA3* (G79-Y337), *ScADG3* (G48-D318), *SpADG3* (T70-D335), *AfSUN1* (T138-Y414), *AfADG3* (G51-E315), *TsSUN1* (G154-Y433), *BcSUN1* (S189-S471) or *BcADG3* (G60-A322), respectively. Plasmids BHUM4052, BHUM4054 and BHM4056 were generated by *in vivo* cloning in *S. cerevisiae* following the same strategy used for plasmid BHUM3446, but using synthetic DNA fragments (Twist Bioscience, San Francisco, CA, USA) for the SUN domain of *ScSUN4* carrying the appropriate mutations.

### Quantitative Cell Cluster Analysis (QCA)

For <u>Q</u>uantitative Cell <u>C</u>luster <u>A</u>nalysis (QCA; Figure S4), diploid yeast strains were grown in liquid YNB medium with appropriate supplements at 30°C into early stationary phase, followed by vigorous stirring of cultures on a vortex shaker. Images of cells and cell clusters were then obtained by low magnification (200x) brightfield microscopy using a Stemi2000-C microscope (Zeiss, Jena, Germany) and digital photography with a Canon EOS1300D CMOS camera. Routinely, twenty images were taken for each yeast strain analyzed. Digital raw images were further processed by the *ImageJ* software program (75) for background correction, cluster segmentation and determination of individual cell cluster areas in µm^2^. Routinely, the sizes of more than 10^3^ random particles were determined for each strain, followed by calculation of the mean cell cluster area using log_10_-transformed data. For statistical analysis and visualization of the data, the *Microsoft Excel* and *R* (www.r-project.org) software programs were used. For each strain, at least three independent QCA measurements were performed. Detailed scripts for image processing and data evaluation are available upon request.

### Production and purification of recombinant SUN domains

For production and purification of the different SUN domains, the *E. coli* SHuffle^®^ T7 Express strain (New England Biolabs, Frankfurt, Germany) carrying appropriate pET-28a-based expression plasmids (Table S8) was used. Bacterial strains were grown at 12°C for 72 h in lysogeny broth (LB) before addition of 0.05 mM IPTG to induce heterologous expression of the SUN domains following a low temperature protocol (76). Cell pellets were then harvested by centrifugation and resuspended in purification buffer (20 mM Tris/HCl, 200 mM NaCl, pH 8.0). Cell disruption was performed mechanically in purification buffer in the presence of EDTA, Lysozyme and PMSF using a French pressure cell press. After centrifugation, the sterile filtered supernatant was loaded onto a Ni-NTA column (Macherey-Nagel, Düren, Germany) and eluted with 50 mM imidazole. Pooled fractions were concentrated with an Amicon Ultra-15 filter unit (EMD Millipore, 10 kDa). Finally, a size exclusion chromatography polishing step was performed on a HiLoad 16/60 Superdex 200 pg column (GE Healthcare, Solingen, Germany) yielding >95% pure recombinant proteins. Routinely, protein yields of 10-15 mg per liter expression culture could be obtained.

### Crystallization of recombinant SUN domains

Initial crystallization attempts were made with the JCSG Core Suites I-IV (Qiagen, Hilden, Germany) using the sitting-drop-vapour diffusion-method in Innovaplate SD-2 96-well plates (Jena Bioscience). 300 nl protein and precipitant drops were set up with a Cartesian 4004 robot system (Marburg Crystallisation Lab, MarXtal). Well diffracting crystals of ScSun4 SUN domains were obtained at 4 °C using JCSG Core Suite II conditions 26 (10% PEG 400, 0.1 M NaHEPES pH 7.5, 1.5 M ammonium sulphate) and 75 (40% PEG 300, 0.1 M MES pH 5.2, 0.2 M MgCl_2_) with ∼50 mg/ml freshly purified protein within 48 h. After soaking with 50 mM GdAc_3_ for 48 h, crystals were flash-frozen in liquid nitrogen. Well diffracting crystals of the SUN domain of ScSim1 were obtained at 5°C within 48 h in a hanging drop scale in optimized conditions (0.1 M MgCl_2_, 0.1 M Tris-HCl pH 7.5, 30 % PEG 4000) using freshly purified protein at a concentration of ∼20 mg/ml.

### Structure determination

For the SUN domain of ScSun4, single-wavelength anomalous diffraction (SAD) data of a gadolinium derivative were collected in-house at 100 K using the FR591 rotating anode X-ray generator (Bruker *AXS*), whereas native datasets were collected using beamline ID14-4 of the European Synchrotron Radiation Facility (Grenoble, France) and beamline MX14.1 of the Helmholtz-Zentrum Berlin, Bessy II. The SUN domain of ScSun4 was crystallized in space group *I*222 with unit cell parameters *a*=62.4 Å, *b*=99.2 Å, *c*=102.2 Å and *α*=β=γ=90° (Table S2). Data reduction was performed with XDS and XSCALE (77). Structure solution by SAD phasing was performed with the structure-determination platform *Auto*-Rickshaw (78). *Phaser* (79) was used for heavy atom search and *MLPhare* (80) for phase refinement. *Pirate* (81) modified the initial electron density with the result that *SHELXE* (82) started to build up the first polyalanine model. After this step, *Resolve* (83) adapted the ScSun4 primary sequence to the polyalanine model. Semiautomatic tracing and model building were done with *ARP/wARP* (84). Final model building and refinement was done with *refmac5* (85) of the CCP4 suite (80) and WinCoot (86) at 1.1 Å resolution and led to 14.1% for R_work_ and 15.7% for *R_free_*, respectively. Likewise, the structure of the SUN domain of ScSim1 was solved and refined at 1.2 Å resolution after molecular replacement using the ScSun4 SUN domain and *Phaser* of the phenix suite.

### Surface plasmon resonance assays (SPR)

The biomolecular interaction of the recombinant SUN domain of ScSun4 towards β-1,3-glucans varying in dense of polymerisation (DP) was measured with a Biacore^TM^ T100 biosensor instrument (GE Healthcare, Solingen, Germany). pH scouting was initially performed varying the buffer pH value to optimize the recombinant protein immobilization. Recombinant ScSun4 SUN domain (0.3 mg/mL in 10 mM acetate buffer, pH 5.0) was then coupled to the surface of a Sensor CM5 Chip (Biacore Inc.) by standard amine chemistry (N-hydroxysuccinimide-1-ethyl-3-[3-dimethylaminopropyl]-carbodiimide to a level of ∼1000 response units. Free active sites were then blocked by 1 M ethanolamine. A reference flow cell with an activated and blocked surface, but without the recombinant protein, was created to normalize readings. Runs were performed with Laminarin (Biosynth Carbosynth, Billingham, UK) (50 mM acetate buffer pH 5.5) varying in dense of polymerisation (DP) for 600 s at 20°C.

### Molecular dynamics simulations

Molecular dynamics analysis of triple helical β-glucan (curdlan) binding by the SUN domain of ScSun4 was performed using AMBER14 (87) with the ff99sb and GLYCAM06 (88) force fields. The SUN domain was positioned in a periodic, water filled and neutralized box. The box size was chosen as 86 x 75 x 91 Å and the molecular dynamics simulations were performed with a step size of 2 fs at 300 K, using an isothermal-isobaric ensemble (NPT). After minimization and equilibration for 2 ns, trajectories were collected for a further 100 ns. Molecular graphics were then generated and analyzed with Visual Molecular Dynamics (VMD1.9.2) (89).

### Bioinformatic and statistical analysis

The SSN analysis based on the Enzyme Similarity Tool of the Enzyme Function Initiative (EFI-EST) (33) was performed on IPR005556, by using an initial BLAST-derived E-value stringency of 10^-5^. In later steps, the alignment score E-value was limited to 10^-60^ and sequence length was restricted to 300-700 amino acids. The resulting network had a pair-wise sequence identity greater than 60 % for each of the 1602 sequences. The data were further analyzed with Cytoscape (90), Clustal Omega (91) and WEBLOGO (92). Figures of protein structures were generated with the Molecular Graphics Software PyMOL v2.3.0 (Schrödinger, LLC). The alignment of protein sequences was performed using Clustal Omega (91). Phylogenetic analysis of protein sequences was performed using T-Coffee (93) and MEGA6 (94) using the maximum-likelihood method. Statistical analysis of data obtained by QCA was performed using the *Microsoft Excel* and *R* (www.r-project.org) software programs. De novo molecular models of SUN paralogs alone or in complex with β-1,3-glucans were done with Alphafold 3 (95) and Chai-1 (96).

## Data availability

The atomic coordinates for the crystal structures of the SUN domains of ScSun4, ScSun4^E283A^, ScSun4^Q318A^ and ScSim1 have been deposited in the Protein Data Bank under accession numbers 9T47, 9T4O, 9T4N and 9T4Q, respectively.

## Acknowledgments

The authors thank Daniela Störmer and Petra Gnau for technical support. We thank the staff of beamline ID14-4 of the European Synchrotron Radiation Facility (Grenoble, France) and of the beamline MX14.1 of the Helmholtz-Zentrum Berlin, Bessy II, Germany, for excellent support. This research was funded by the German Research Foundation (DFG) with grants to H.-U.M. and L.-O.E. from CRC 987.

